# A Systematic Review and Meta-analysis for Association of *Helicobacter pylori* Colonization and Celiac Disease

**DOI:** 10.1101/2020.10.12.335836

**Authors:** Fazel Isapanah Amlashi, Zahra Norouzi, Ahmad Sohrabi, Hesamaddin Shirzad-Aski, Alireza Norouzi, Ali Ashkbari, Naghme Gilani, Seyed Alireza Fatemi, Sima Besharat

## Abstract

**Background and objectives:** Based on some previous observational studies there is a theory that suggests a potential relationship between *Helicobacter pylori* (*H. pylori*) colonization and celiac disease (CD), however, the type of this relationship is still controversial. Therefore, we aimed to conduct a systematic review and meta-analysis to explore all related primary studies to find any possible association between CD and human *H. pylori* colonization.

**Data sources:** Studies were systematically searched and collected from four databases and different types of gray literature to cover all available evidence. After screening, the quality and risk of bias assessment of the selected articles were evaluated.

**Synthesis methods:** Meta-analysis calculated pooled odds ratio (OR) on the extracted data. Furthermore, heterogeneity, sensitivity, subgroups, and publication bias analyses were assessed.

**Results:** Twenty-four studies were included in this systematic review, with a total of 5241 cases and 132947 control people. The results of meta-analysis on 24 studies showed a significant and negative association between *H. pylori* colonization and CD (pooled OR= 0.58; 95% CI = 0.45 - 0.76; P < 0.001), with no publication bias (P = 0.407). The L’Abbé plots also showed a trend of having more *H. pylori* colonization in the control group. Among subgroups, ORs were notably different only when the data were stratified by continents or risk of bias; however, subgroup analysis could not determine the source of heterogeneity.

**Conclusions:** According to the meta-analysis, *H. pylori* has a mild protective role toward CD. Although this negative association is not strong, it is statistically significant and should be further considered. Further investigations in both molecular and clinic fields with proper methodology and more detailed information are needed to discover more evidence and underlying mechanisms to clear the interactive aspects of *H. pylori* colonization in CD patients.

**Systematic review registration number (PROSPERO):** CRD42020167730

## Introduction

Celiac disease (CD) is an inflammatory autoimmune disorder that occurs in genetically predisposed persons who develop an immune reaction to gluten [1]. CD is a multifactorial disease that has risen from the interaction between gluten and immune, genetic, and environmental factors and can be diagnosed at any age [2]. The global prevalence of CD is around 1% [3]. In this regard, the high prevalence of CD was firstly observed in Sweden and Finland but now many studies from Europe reported the prevalence of CD to be around 1% or even higher [4]. Furthermore, population-based studies from Asia showed up to 1.3% for the prevalence of CD [5]. So generally, the incidence of CD is considered to be increasing in many countries [6–8].

There are three models of CD manifestations including individuals who show no symptoms at all, patients with non-specific manifestations of nutritional deficiency, and patients who present the severe form of the CD [3]. Although CD affects the small intestine, clinical manifestations are wide, and we can perceive extra-intestinal symptoms, including anemia, raised transaminases, recurrent miscarriages, osteoporosis, associated autoimmune disorders, and aphthous stomatitis. Primary manifestations can be diarrhea and weight loss along with atypical symptoms such as constipation and recurrent abdominal pain [9,10]. The only effective therapy of CD is a gluten-free diet, as the small intestine is sensitive to gluten-rich products in these people. Persistent or recurrent symptoms occur when they do not follow properly a gluten-free diet. Patients who have poor control over a gluten-free diet may experience complications such as refractory sprue, small intestinal adenocarcinoma, and enteropathy-associated T-cell lymphoma [3]. There are some diagnostic methods for CD detection, such as serology tests, which are the most important and accessible one and can identify CD at early stages, and include anti-tissue transglutaminase IgA antibodies (anti-tTG-IgA), IgG anti-deamidated gliadin peptide antibodies (DPG IgG), and anti-endomysial antibody (EMA) tests. Moreover, molecular methods such as detection of HLA-DQ2 and HLA-DQ8 can help the diagnosis of gluten sensitivity [10]. *H. pylori* is one of the most common chronic bacterial infections worldwide, it can cause significant gastroduodenal diseases [11]. The infection affects up to 90% of the population in developing countries and its prevalence is less than 40% in developed countries. Generally, half of the global population can have *H. pylori* colonization [12]. Different methods are available for the detection of *H. pylori* including culture, histology, rapid urease test (RUT), polymerase chain reaction (PCR), serology, stool antigen test, and urea breath test (UBT) [11]. Epidemiological and histomorphological studies show that *H. pylori* is an important etiological factor in patients with peptic ulcer disease, primary gastric mucosa associated lymphoid tissue (MALT) lymphoma, and gastric cancer [13]. Besides, it is one of the rare bacteria that can be involved in many autoimmune diseases. Several studies focused on the probable relation between *H. pylori* colonization and several autoimmune diseases such as multiple sclerosis, autoimmune thyroid disease, immune thrombocytopenic purpura, and neuromyelitis optica [14]. *H. pylori* can cause microscopic duodenal inflammation and can be related to more severe damage associated with CD [15]. In genetically predisposed persons, *H. pylori* infection can influence immune responses in the small intestine and may advance to gluten-related enteropathy [16–18]. A study at Colombia University showed increased intraepithelial lymphocytes in the duodenal mucosa in patients with *H. pylori* infection that could be reversed by the eradication of *H. pylori* [19]. Some studies reported no relationship between *H. pylori* and CD [20–22]; whereas, others stated that *H. pylori* can protect against CD [15,23].

Up to now and based on our search, no systematic review and meta-analysis have been conducted about this subject. Due to the inconsistency of the previous primary studies, we aimed to prepare a systematic review to scrutinize any possible association between *H. pylori* colonization and CD.

## Materials and Methods

### Eligibility Criteria

The design of the present study was based on the methodology of the Preferred Reporting Items for Systematic Reviews and Meta-Analyses (PRISMA) guideline [24]. This review is registered in the International prospective register of systematic reviews (PROSPERO) with registration number CRD42020167730 (Available: https://www.crd.york.ac.uk/prospero/display_record.php?RecordID=167730).

### Inclusion/exclusion criteria

Inclusion criteria included: 1) Case-control, cross-sectional, and brief-report studies that contain evidence about the relationship between *H. pylori* and CD, 2) Human clinical study, 3) Having at least one parameter of the relative risk effect sizes such as risk ratio (RR), odds ratio (OR), and hazard ratio (HR) or can be calculated based on the information of *H. pylori* status in both case and control groups.

The status of *H. pylori* should be detected by UBT, RUT, culture, enzyme-linked immunosorbent assay (ELISA), histology, immunohistochemistry (IHC), and PCR methods. Furthermore, the CD should be confirmed by endoscopy and biopsy ± serology tests and/or HLA DQ2/DQ8 genotyping. Articles without any of this information were excluded from further analysis.

### Sources and search strategy

All relevant articles, published from 1990 to the end of 2019 were collected from the electronic database of PubMed, ProQuest, Scopus, and Web of Science. The ProQuest database was also reviewed for related dissertations, manually. Other related protocols were manually searched in PROSPERO. Besides, to expand the scope of the search and find any additional studies, we reviewed the different types of grey literature such as meeting and conference abstracts for relevant articles (including Digestive Disease Week^®^ (DDW), United European Gastroenterology (UEG), and American College of Gastroenterology). Moreover, we searched manually some key journals in the CD and *H. pylori* fields, including “World Journal of Gastroenterology”, “Journal of Pediatric Gastroenterology and Nutrition”, “Gut”, “Gastroenterology”, and “Helicobacter”. In the end, we performed a manual search in the references of the selected articles. We did not impose geographical or linguistic restrictions on our search, and non-English studies with English abstracts were also included.

The Medical subject heading (MeSH) database was used to find various terms of CD and *H. pylori*. Two main keywords were “*Helicobacter pylori*” and “Celiac”. For better searching in the databases, we made syntaxes from a combination of free-text method, MeSH terms, the keywords, and Boolean operators (AND/OR/NOT). Moreover, calculating NNR (number need to read) helped us in evaluating the output of syntax and dedicating. The following syntax was applied in PubMed and adjusted for each search engine based on its search guidelines:

(Celiac OR “Celiac disease” OR Coeliac OR “Coeliac disease” OR (Disease AND Celiac) OR (Disease AND Coeliac) OR “Gluten Enteropathy” OR (Enteropathy AND Gluten) OR “Gluten Enteropathies” OR (Enteropathies AND Gluten) OR “Gluten-Sensitive Enteropathy” OR (Enteropathy AND Gluten-Sensitive) OR “Gluten-Sensitive Enteropathies” OR (Enteropathies AND Gluten-Sensitive) OR (Sprue AND Celiac) OR (Sprue AND Coeliac) OR (Sprue AND Nontropical) OR “Nontropical Sprue” OR “Celiac Sprue” OR “Coeliac Sprue” OR “Sprue” “Endemic Sprue” OR “Gluten Intolerance”) AND (“*Helicobacter pylori*” OR “*Campylobacter pylori*” OR “*Campylobacter pyloridis*” OR “*Campylobacter pyloris*” OR “*Helicobacter nemestrinae*” OR “*Helicobacter* infection” OR “HP infection” OR “HP infections” OR “*Helicobacter* infections” OR “*Helicobacter* colonization” OR “HP colonization” OR “*Helicobacter* colonization” OR “*H Pylori*” OR “enterohepatic *Helicobacter*” OR “EHS” OR “*Helicobacter*”) AND 1990/01/01:2019/12/31[dp]

### Quality and risk of bias assessment

The search outputs were exported into the Endnote software (Version X7; Thompson Reuters Corporation, Toronto, ON, Canada) to remove any duplicate and screen articles. The first step of screening for determining the eligible primary articles was done based on the titles and abstracts. Two independent reviewers (A.A. and A.F.) evaluated the selected full-text articles and separately classified them into three relevant, irrelevant, and unsure groups, based on the eligibility criteria. Any discrepancy was supervised by a third reviewer (S.B.) and resolved via a consensus. If there was still a doubt in the selection of a study, the whole team made the final decision.

The quality and risk of bias assessment of the selected articles were independently evaluated by two reviewers (Z.N. and N.G.), using a modified checklist of the Newcastle-Ottawa Scale (NOS) form for case-control studies [25]. The studies were classified into three poor, fair, and good categories based on getting a score between zero and eight in the selection, comparability, and outcome/exposure domains. Consensus and the opinion of the third reviewer (S.B) resolved any disagreements between the two reviewers.

### Data extraction

Two authors (Z.N, F.A) independently performed data extraction from each paper based on a defined protocol. The data were divided into three categories and included 1) General information: the first author’s names, year of publication, journal names, country, and region; 2) The risk of bias assessment; 3) Study setting: study design, study duration, sample size and total population recruited in each study, inclusion and exclusion criteria, age group and age range, sex, the definition of CD, source of the data, diagnostic methods, OR and 95% CI, and the ethical approval. In cases of any missing or additional necessary data, we further sent an e-mail to each author, in charge of the related study.

### Data synthesis and analysis

We used the R software environment, version 4.0.2, and the “meta” package to calculate pooled OR and 95% CIs for each study. Based on the methodological heterogeneity between the studies, the random-effect model (REM) was used for the combination method. The size of the combined effect was calculated and displayed in a forest plot. The standard chi-square test (*Q* Cochrane test) and *I^2^* scale evaluated the heterogeneity between studies. Combinations with *I^2^* scale equal or more than 70% assumed as a severe heterogeneity. In addition, Drapery plot was drawn to show the P-value functions for the included studies, as well as pooled estimates and to provide two-sided confidence intervals for all possible alpha levels (confidence interval function) [26,27]. For finding the cause(s) of heterogeneity, subgroup and sensitivity analysis was used. The studies were stratified based on each factor, including the median of publication year, overall age group of the participants, continent and regions of the studies, *H. pylori* detection method, the risk of bias assessment, as well as sampling quality. In the last subgroup, sampling, the articles were categorized into two groups (I) “Appropriate sampling” or (II) “Not appropriate sampling” groups. The studies were located in the “Not appropriate sampling” group, if the number of participants in each case and control group was less or equal than 50 people or the ratio of control to the cases was more than four times or less than 90%.

The Funnel chart, followed by Begg’s test analyzed possible publication bias. Baujat and L’Abbé plots were used to show the effect of each study on the heterogeneity and overall influence of the results [28–30]. Finally, a one-out remove method was used to detect the influence of each study on the overall pooled OR.

## Results

### Search results

In total, the general searching step obtained 1641 papers. After removing duplicates, 1050 studies were screened, based on their titles and abstracts. Among them, 963 articles were excluded due to not meeting the inclusion criteria. Therefore, the full-texts of 87 remained articles were completely evaluated. Finally, 24 studies were included in the systematic review and also meta-analysis (64% agreement between reviewers; κ=0.83) [15,16,34–43,18,44–47,20–23,31–33], and the others were excluded due to a reason mentioned in Fig 1.

**Fig 1.**
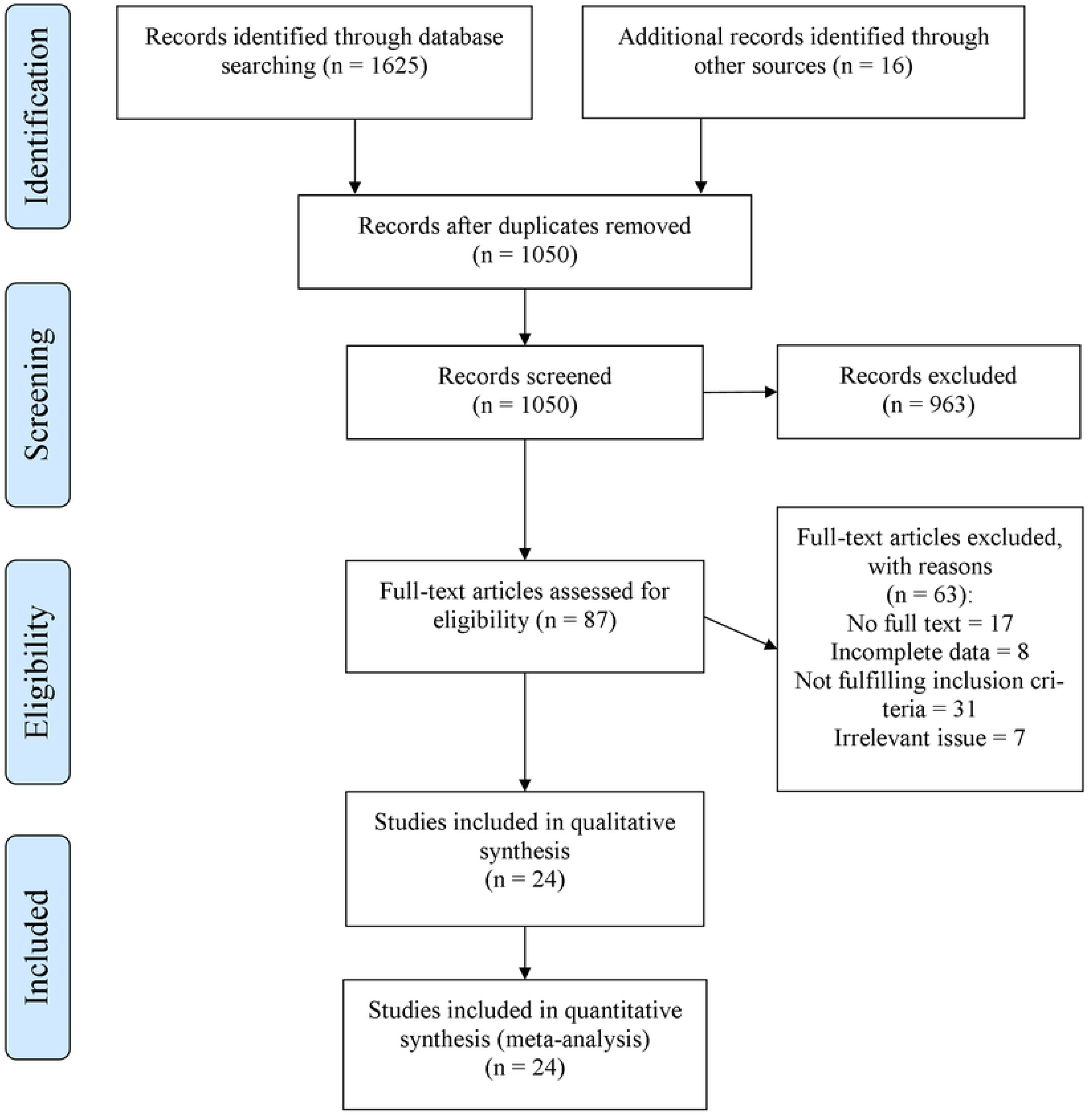
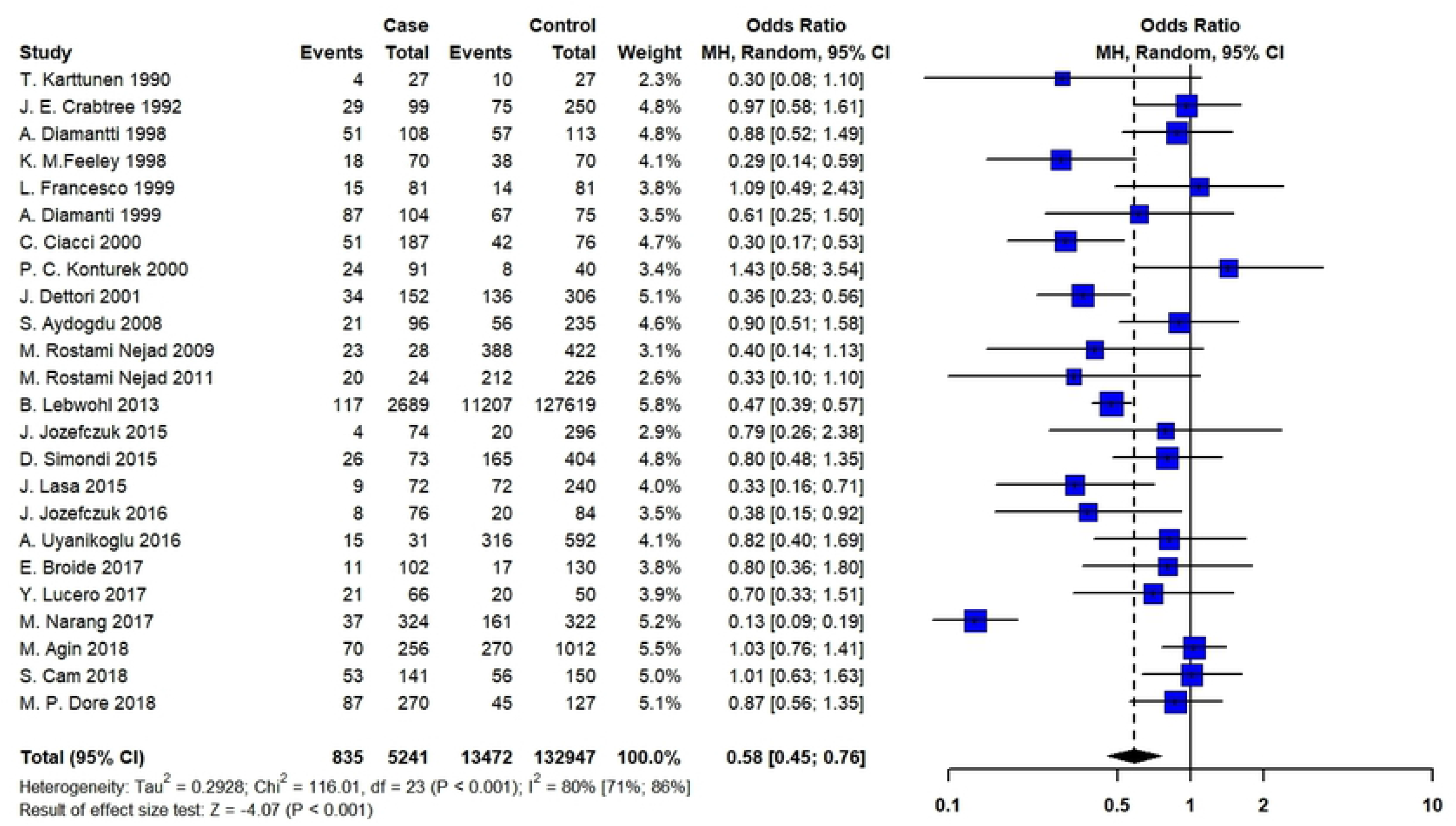
Flow diagram of literature search, screening, and selection of studies for review and metaanalysis.

### Characteristics of the included articles

The designs of all the included studies were case-control, with a total of 5241 cases and 132947 control people. Among the studies, Lebwohl *et al*. had the highest study population with 130308 participants (included 2689 cases and 127619 controls) [15]. Most studies were from Italy (N = 5) [16,36,38,42,45] and Turkey (N = 4) [20–23]. Some studies had been performed only on children [18,20–22,36,40,43] or adults [23,31,35,37,38,42,45], the others evaluated both age groups [15,16,34,46,47]. Table 1 shows the details of the included articles. The NOS assessment showed that only 20.8% of the studies had good quality.

**Table 1.** Main characteristics of the included studies in the meta-analysis on the *Helicobacter pylori* infection and celiac disease (CD)

| First author | Year | Journal title | Country | Definition of CD | Risk of bias assessment | Case-number | Control - number | Laboratory tests of *H. pylori* |
| --- | --- | --- | --- | --- | --- | --- | --- | --- |
| Karttunen | 1990 | Journal of Clinical Pathology | Finland | Registry | F<sup>c</sup> | 27 | 27 | IHC<sup>f</sup> |
| Crabtree | 1992 | Journal of Clinical Pathology | England | Endoscopy + Serology + Positive response to GFD<sup>b</sup> | P<sup>d</sup> | 99 | 250 | ELISA<sup>g</sup> |
| Feeley | 1998 | Journal of Clinical Pathology | Ireland | Registry | F | 70 | 70 | IHC |
| Diamantti | 1998 | Intestinal Disorders (poster) | Argentina | Endoscopy | P | 108 | 113 | IHC |
| Diamantti | 1999 | The American Journal of Gastroenterology | Argentina | Endoscopy + Serology + Positive response to GFD | P | 104 | 75 | IHC |
| Francesco | 1999 | Journal of Pediatric Gastroenterology and Nutrition | Italy | Endoscopy+ Positive response to GFD | G<sup>e</sup> | 81 | 81 | IHC + RUT<sup>h</sup> + ELISA |
| Konturek | 2000 | The American Journal of Gastroenterology | Germany | Endoscopy + Serology | F | 91 | 40 | ELISA |
| Ciacchi | 2000 | European Journal of Gastroenterology and Hepatology | Italy | Endoscopy | P | 187 | 76 | IHC + ELISA |
| Dettori | 2001 | Digestive and Liver Disease (Poster) | Italy | Endoscopy + Serology | G | 152 | 306 | UBT<sup>i</sup> + IHC |
| Aydogdu | 2008 | Scandinavian Journal of Gastroenterology | Turkey | Endoscopy + Serology | P | 96 | 235 | IHC |
| Rostami-Nejad | 2009 | Revista espanola de enfermedades digestivas | Iran | Endoscopy + Serology | P | 28 | 422 | IHC |
| Rostami-Nejad | 2011 | Archives of Iranian Medicine | Iran | Endoscopy + Serology | P | 24 | 226 | IHC |
| Lebwohl | 2013 | American Journal of Epidemiology | American | Endoscopy | F | 2689 | 127619 | IHC |
| Simondi | 2015 | Clinics and Research in Hepatology and Gastroenterology | Italy | Endoscopy + Serology + HLA DQ2/DQ8 genotyping | F | 73 | 404 | UBT + IHC |
| Jozefczuk | 2015 | European Review for Medical and Pharmacological Science | Poland | Endoscopy + Serology | F | 74 | 296 | UBT |
| Lasa | 2015 | Archivos de Gastroenterologia | Argentina | Endoscopy + Serology | G | 72 | 240 | IHC |
| Jozefczuk | 2016 | European Review for Medical and Pharmacological Science | Poland | Endoscopy + Serology | F | 76 | 84 | UBT + ELISA |
| Uyanikoglu | 2016 | Euroasian Journal of Hepato-Gastroenterology | Turkey | Endoscopy | F | 31 | 592 | IHC |
| Narang | 2017 | Journal of Gastroenterology and Hepatology | India | Endoscopy + Serology + Positive response to GFD | G | 324 | 322 | IHC + RUT |
| Broide | 2017 | AGA abstracts | Germany | Endoscopy + Serology | F | 50 | 50 | UBT + IHC |
| Lucero | 2017 | Frontiers in Cellular and Infection Microbiology | Chile | Endoscopy + Serology + HLA DQ2/DQ8 genotyping | G | 66 | 50 | IHC + RUT |
| Agin | 2018 | Archives of Medical Science | Turkey | Endoscopy + Serology | P | 256 | 1012 | IHC |
| Cam | 2018 | EHMSG <sup>a</sup> | Turkey | Endoscopy | P | 141 | 150 | UBT + IHC |
| Dore | 2018 | Helicobacter | Italy | Endoscopy + HLA DQ2/DQ8 genotyping | F | 270 | 127 | UBT + IHC |

<sup>a</sup>XXXIst International Workshop on *Helicobacter* & Microbiota in Inflammation & Cancer; <sup>b</sup>gluten-free diet; <sup>c</sup>Fair; <sup>d</sup>Poor; <sup>e</sup>Good; <sup>f</sup>Immunohistochemistry; <sup>g</sup>Enzyme-linked immunosorbent assay; <sup>h</sup>Rapid Urease Test; <sup>i</sup>Urease Breath Test.

### Association between *H. pylori* and CD

By analyzing a combination of 24 studies that addressed the association between *H. pylori* colonization and CD, a negative association was found (pooled OR = 0.58; 95% CI = 0.45 – 0.76; P ≤ 0.001). The REM and Chi-squared method showed a significant statistical heterogeneity between the studies I^2^ = 80%, P ≤ 0.001, and X^2^ = 116, P ≤ 0.001, respectively (Fig 2).

**Fig 2.**
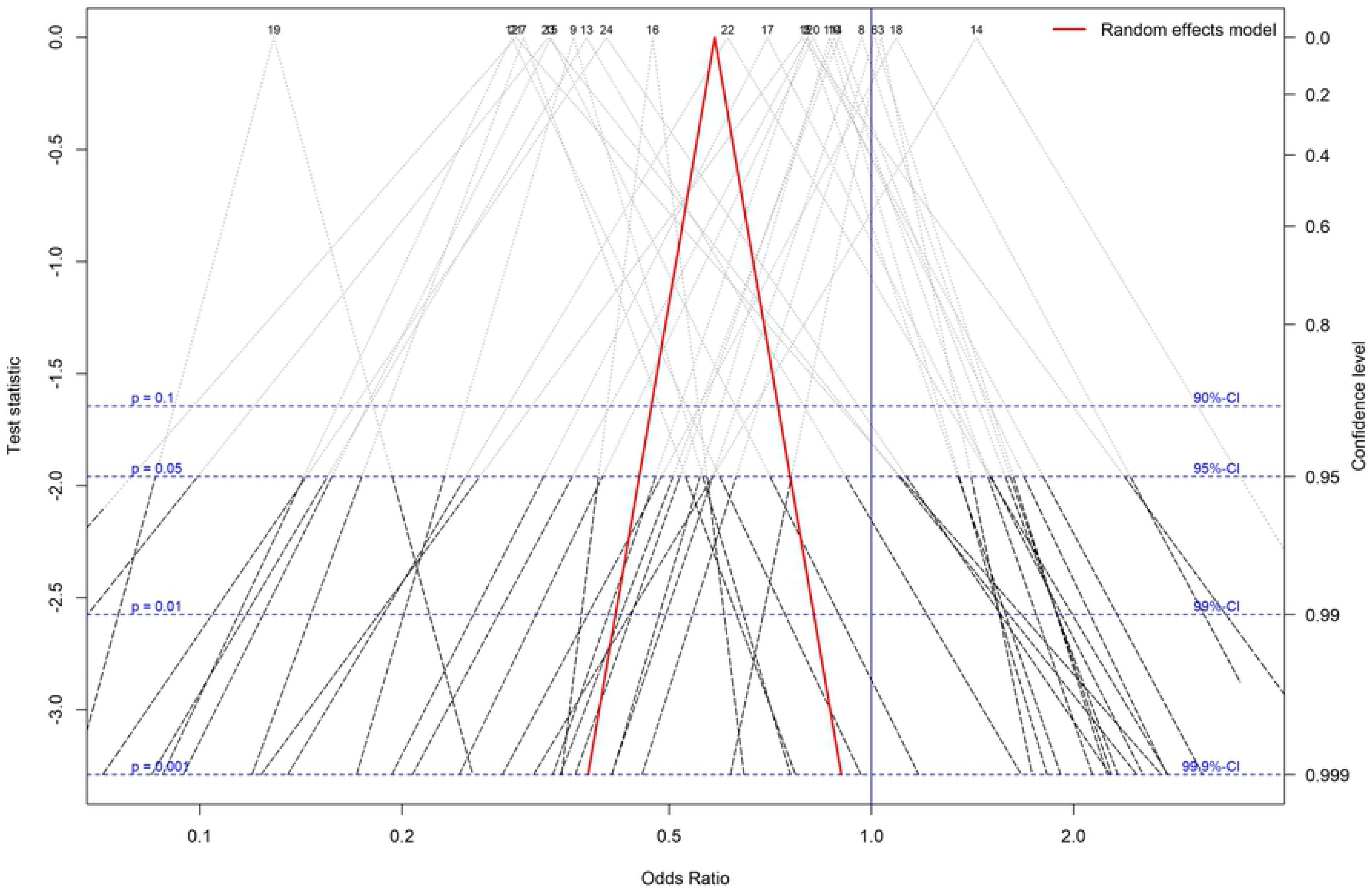
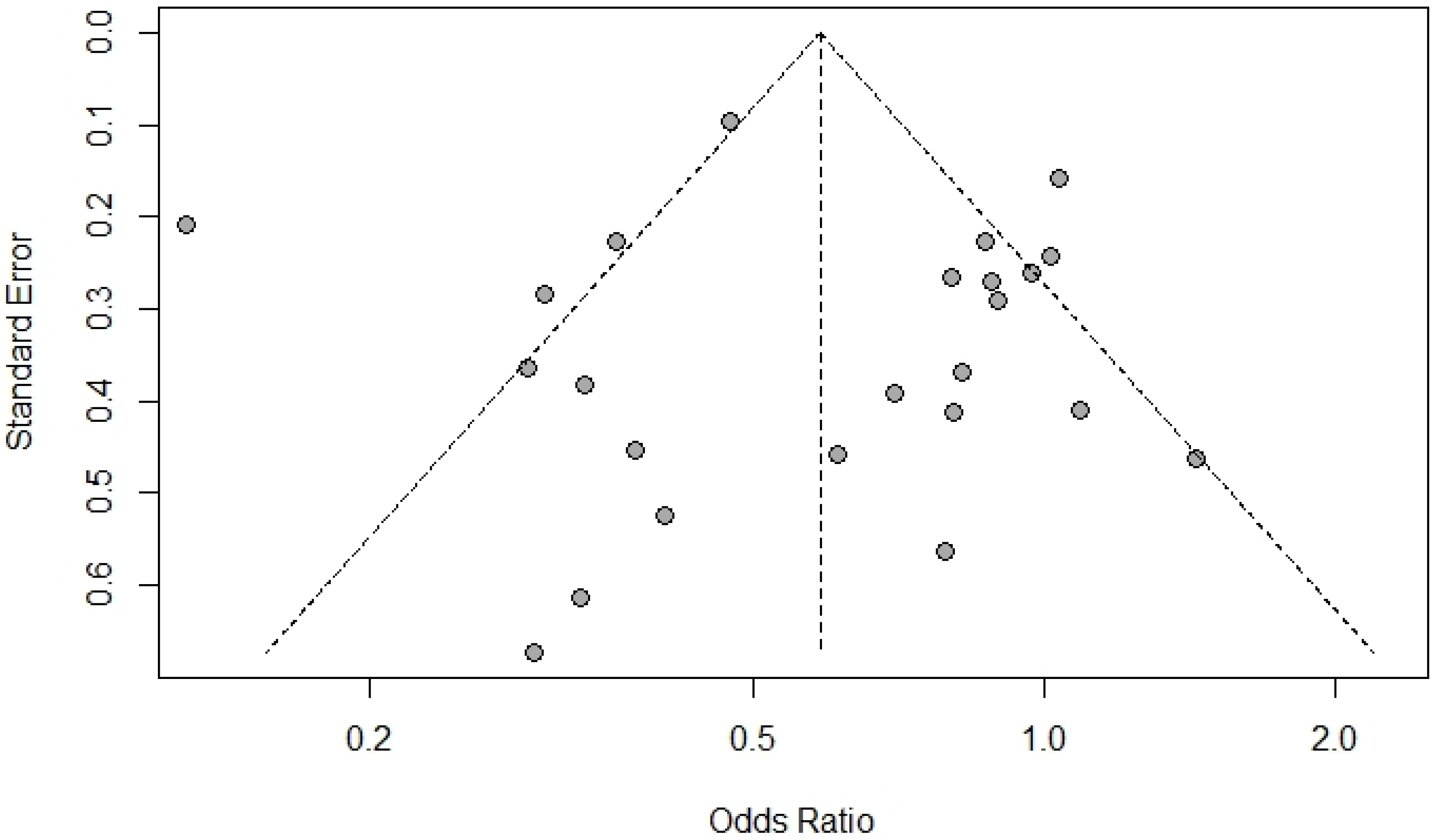
The Forest plot of the meta-analysis on the association of the *Helicobacter pylori* and celiac disease. The random-effects model was used to calculate pooled ORs with 95% CI. CI, Confidence interval; OR, Odds ratio.

Fig 3 (A) shows a Drapery plot, presenting that the estimated effect is less than 1 in the majority of studies, similarly as shown in the Funnel plot. As the horizontal dashed lines show the CIs for common alpha levels (0.1, 0.05, 0.01, 0.001), at any level of prediction interval (90, 95, 99, 99.9%), the main triangle line (the red-bolded line in the online version) did not intersect the null effect line (Incidence Rate Ratio (IRR) = 1). According to the results of the Funnel plot and Begg’s tests, there was no significant publication bias (P = 0.407, Fig 2 B). The Baujat plot (Fig 2 C) showed that the study of Narang *et al*. contributes more than the other studies on the overall heterogeneity and influences the pooled effect size. The L’Abbé plots (Fig 2 D) also showed a trend of having more *H. pylori* colonization in the control group.

**Fig 3.**
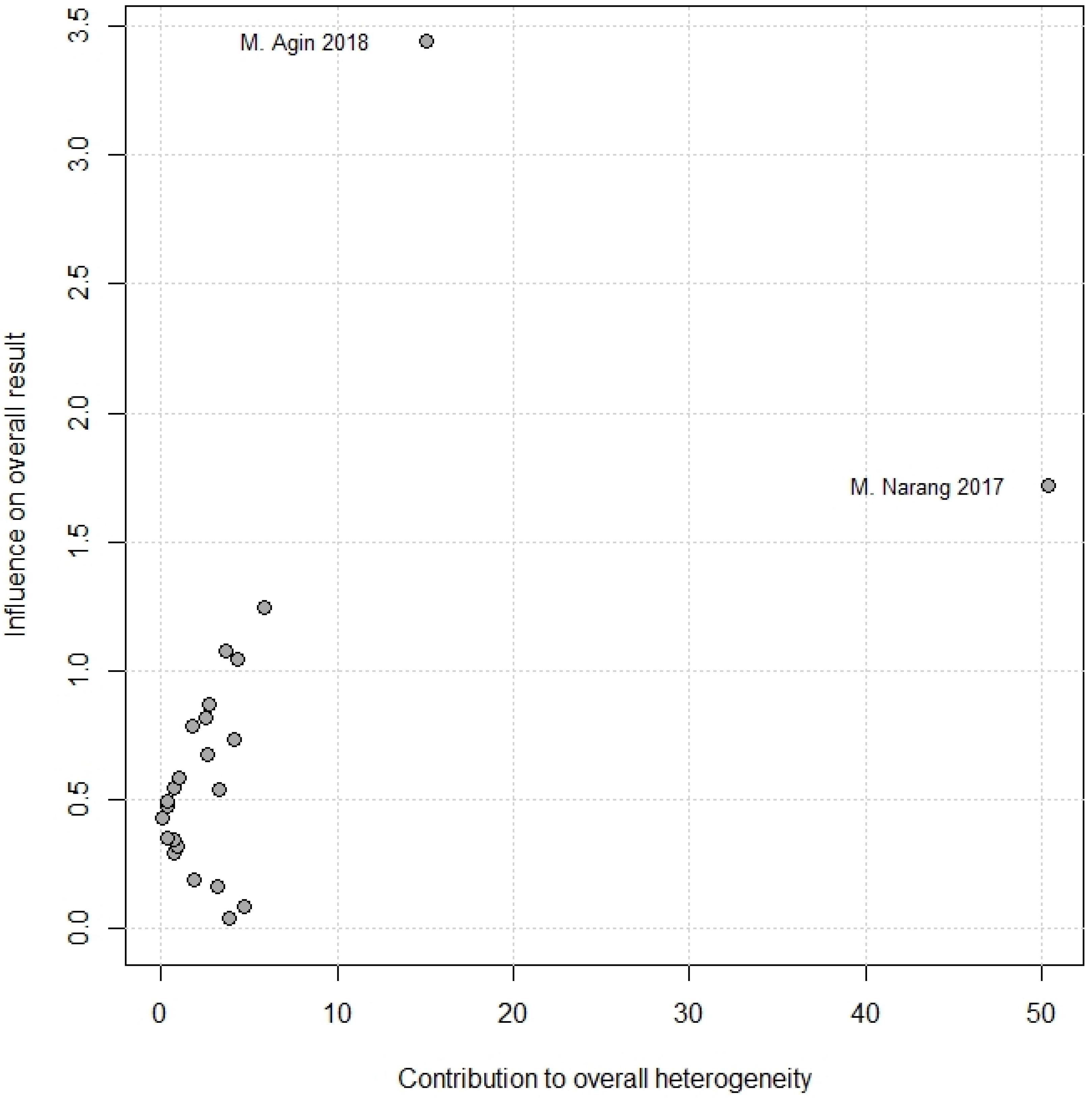
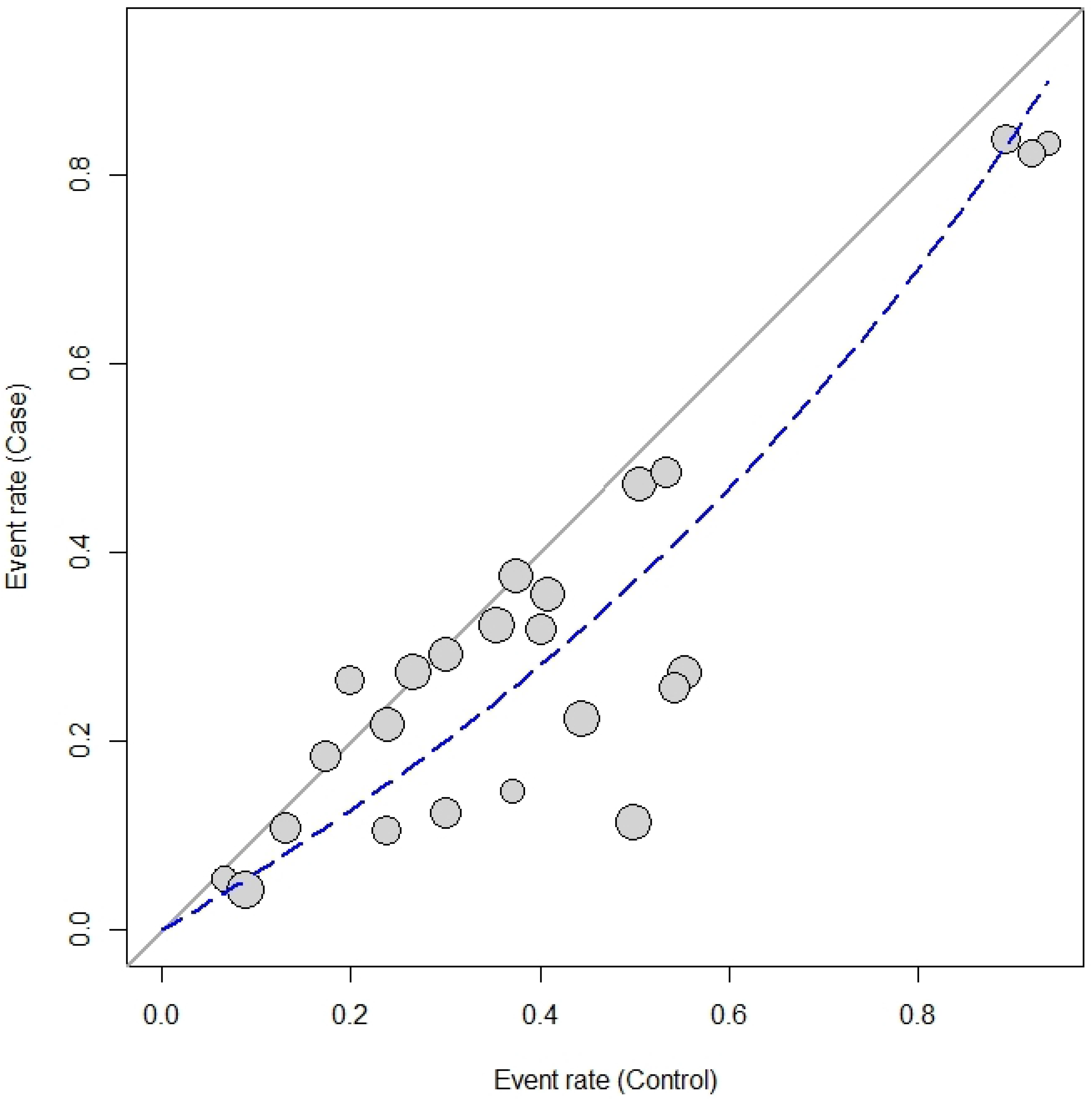
**(A)** The drapery plot shows p-value functions for the included studies as well as pooled estimates and the two-sided confidence intervals for all possible alpha levels (confidence interval function). Each number represents a study. The red-bolded line (in the online version; bold line in the printed version) indicates the range of pooled odds ratio (OR) in each alpha level. **(B)** The Funnel chart shows no significant publication bias, as all studies are in a symmetric scheme. **(C)** The Baujat plot shows the effect of each study on the heterogeneity and overall influence of the results. As can be seen in the graph, the study of Narang et al. contributes more than the other studies to the overall heterogeneity. Each circle represents a study. **(D)** The L’Abbé plot shows more positive results for H. pylori colonization in the control group. The diagonal (x = y) oblique line represents the odds ratio (OR) equal to one. The dashed line indicates the OR of the studies. The size of each circle indicates the assigned random weight of each study. As the picture shows, most studies show an OR of less than one.

### Subgroup and sensitivity analyses

The heterogeneity between studies was evaluated by multiple subgroup analyses in the six subgroups. Based on the continents of the studies, there were few studies in Asia, which made the results unreliable for the statistical analysis [18,46,47]. Europe has a higher OR (OR = 0.69, 95% CI = 0.54 – 0.90) than America (OR = 0.55, 95% CI = 0.40 – 0.76) and Asia (OR = 0.23, 95% CI = 0.10 – 0.52). However, further evaluation is needed due to the low rate of studies, especially in Asia. Other subgroups could not determine the source of heterogeneity (Table 2).

**Table 2.**
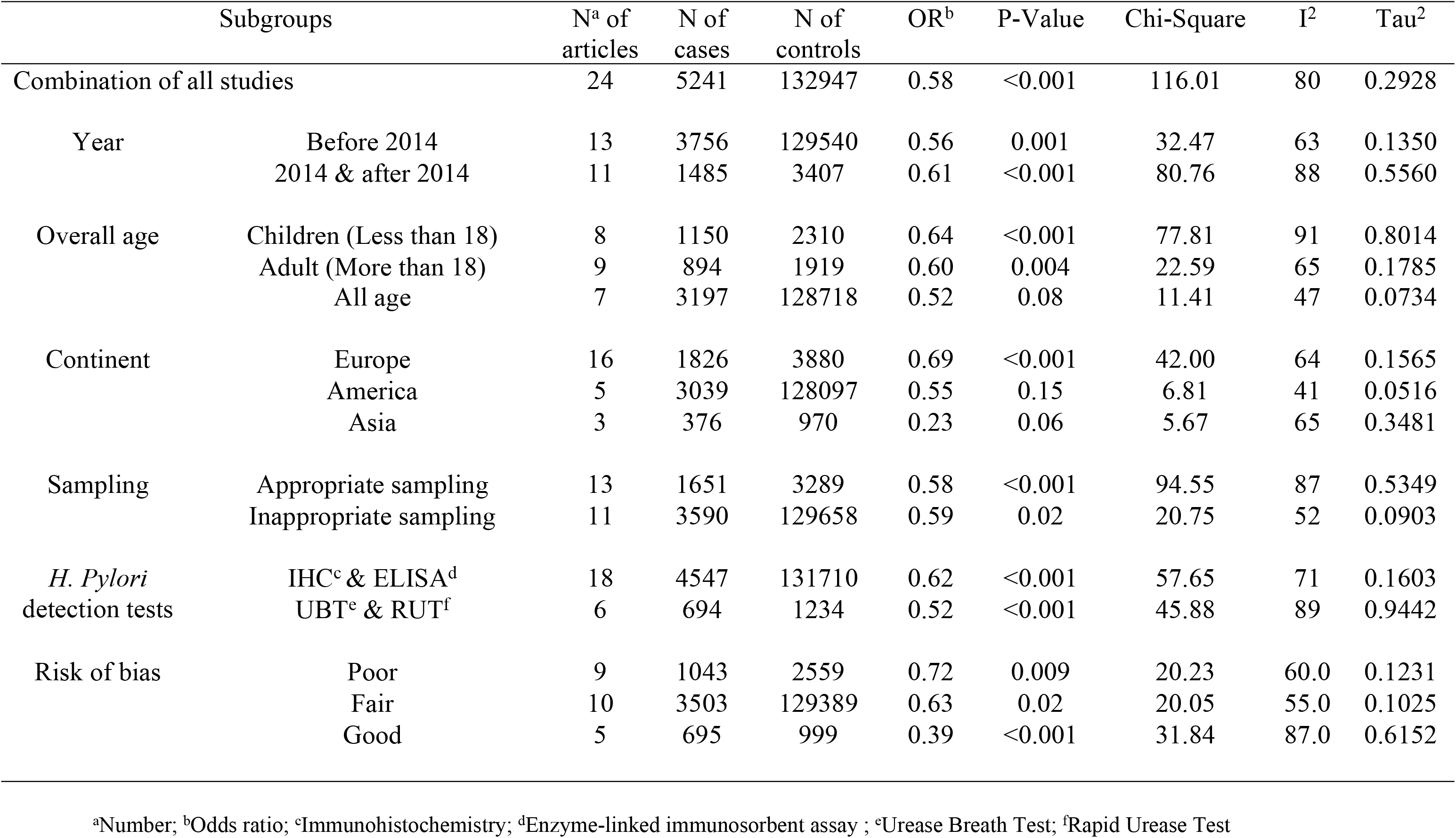
Different subgroup analysis by stratifying the data

| Subgroups |  | N<sup>a</sup> of articles | N of cases | N of controls | OR<sup>b</sup> | P-Value | Chi-Square | I<sup>2</sup> | Tau<sup>2</sup> |
| --- | --- | --- | --- | --- | --- | --- | --- | --- | --- |
| Combination of all studies |  | 24 | 5241 | 132947 | 0.58 | <0.001 | 116.01 | 80 | 0.2928 |
| Year | Before 2014 | 13 | 3756 | 129540 | 0.56 | 0.001 | 32.47 | 63 | 0.1350 |
|  | 2014 & after 2014 | 11 | 1485 | 3407 | 0.61 | <0.001 | 80.76 | 88 | 0.5560 |
| Overall age | Children (Less than 18) | 8 | 1150 | 2310 | 0.64 | <0.001 | 77.81 | 91 | 0.8014 |
|  | Adult (More than 18) | 9 | 894 | 1919 | 0.60 | 0.004 | 22.59 | 65 | 0.1785 |
|  | All age | 7 | 3197 | 128718 | 0.52 | 0.08 | 11.41 | 47 | 0.0734 |
| Continent | Europe | 16 | 1826 | 3880 | 0.69 | <0.001 | 42.00 | 64 | 0.1565 |
|  | America | 5 | 3039 | 128097 | 0.55 | 0.15 | 6.81 | 41 | 0.0516 |
|  | Asia | 3 | 376 | 970 | 0.23 | 0.06 | 5.67 | 65 | 0.3481 |
| Sampling | Appropriate sampling | 13 | 1651 | 3289 | 0.58 | <0.001 | 94.55 | 87 | 0.5349 |
|  | Inappropriate sampling | 11 | 3590 | 129658 | 0.59 | 0.02 | 20.75 | 52 | 0.0903 |
| *H. Pylori* detection tests | IHC<sup>c</sup> & ELISA<sup>d</sup> | 18 | 4547 | 131710 | 0.62 | <0.001 | 57.65 | 71 | 0.1603 |
|  | UBT<sup>e</sup> & RUT<sup>f</sup> | 6 | 694 | 1234 | 0.52 | <0.001 | 45.88 | 89 | 0.9442 |
| Risk of bias | Poor | 9 | 1043 | 2559 | 0.72 | 0.009 | 20.23 | 60.0 | 0.1231 |
|  | Fair | 10 | 3503 | 129389 | 0.63 | 0.02 | 20.05 | 55.0 | 0.1025 |
|  | Good | 5 | 695 | 999 | 0.39 | <0.001 | 31.84 | 87.0 | 0.6152 |

<sup>a</sup>Number; <sup>b</sup>Odds ratio; <sup>c</sup>Immunohistochemistry; <sup>d</sup>Enzyme-linked immunosorbent assay ; <sup>e</sup>Urease Breath Test; <sup>f</sup>Rapid Urease Test

Based on the one-out removed method, the removal of any of the studies could not significantly affect the pooled results. The sensitivity analysis was used to find possible outlier studies with more influence on the results. In this regard, the studies with poor design or not appropriate sampling (the most influential factors in the methodology) were omitted, and the overall effect size and level of heterogeneity were recalculated. The overall OR decreased from 0.58 (95% CI = 0.45 – 0.76) to 0.41 (95% CI: 0.24 – 0.71) but no change in the heterogeneity was observed. The results of the sensitivity analysis confirmed that the correct methodology was significantly associated with a lower amount of overall OR (Table 3).

**Table 3.** Subgroups Sensitivity Analysis

| Models | Subgroups | N<sup>a</sup> of articles | N of cases | N of controls | OR<sup>b</sup> | P-Value | Chi-Square | I<sup>2</sup> | Tau<sup>2</sup> |
| --- | --- | --- | --- | --- | --- | --- | --- | --- | --- |
| All studies |  | 24 | 5241 | 132947 | 0.58 | <0.001 | 116.01 | 80 | 0.2928 |
| Without “poor” quality studies |  | 15 | 4198 | 130388 | 0.53 | <0.001 | 70.87 | 80 | 0.3088 |
|  | Europe | 11 | 1047 | 2157 | **0.64**<sup>†</sup> | 0.02 | 70.87 | 54 | 0.1450 |
|  | America | 3 | 2827 | 127909 | **0.47** | 0.40 | 1.84 | 0 | 0 |
|  | Asia | 1 | 324 | 322 | **0.13** | NA | NA | NA | NA |
|  | Appropriate sampling | 8 | 951 | 1529 | **0.41** | <0.001 | 36.09 | 81 | 0.4663 |
|  | Inappropriate sampling | 7 | 3247 | 128859 | **0.70** | 0.02 | 15.34 | 61 | 0.1076 |
| Without “not appropriate sampling” studies |  | 13 | 1651 | 3289 | 0.58 | <0.001 | 94.55 | 87 | 0.5349 |
|  | Poor | 5 | 700 | 1760 | **0.98** | 0.98 | 0.4 | 0 | 0 |
|  | Fair | 4 | 322 | 580 | **0.49** | 0.21 | 4.5 | 33 | 0.0961 |
|  | Good | 4 | 629 | 949 | **0.34** | <0.001 | 26.14 | 89 | 0.6170 |
|  | Europe | 10 | 1147 | 2614 | **0.71** | 0.001 | 27.46 | 67 | 0.1686 |
|  | America | 2 | 180 | 353 | **0.56** | 0.04 | 4.33 | 77 | 0.3658 |
|  | Asia | 1 | 324 | 322 | **0.13** | NA | NA | NA | NA |
| Without “poor” quality and “not appropriate sampling” studies |  | 8 | 951 | 1529 | 0.41 | <0.001 | 36.09 | 81 | 0.4663 |
|  | Fair | 4 | 322 | 580 | **0.49** | 0.21 | 4.5 | 33 | 0.0961 |
|  | Good | 4 | 629 | 949 | **0.34** | <0.001 | 26.14 | 89 | 0.6170 |
|  | Europe | 6 | 555 | 967 | **0.51** | 0.07 | 10.07 | 50 | 0.1433 |
|  | America | 1 | 72 | 240 | **0.33** | NA | NA | NA | NA |
|  | Asia | 1 | 324 | 322 | **0.13** | NA | NA | NA | NA |

<sup>a</sup>number; <sup>b</sup>odds ratio; <sup>†</sup>The bolded items show a statistical difference in OR of region subgroup.

## Discussion

Most information about the relationship between *H. pylori* colonization and CD is inferred from case-control studies and more prospective cohort and /or systematic review are needed to provide the strongest evidence about this association. In this regard, the present study is the first systematic review and meta-analysis that aimed to clarify any possible association between *H. pylori* colonization and CD. The meta-analysis showed a fair negative relationship between *H. pylori* and CD, and confirmed the findings of some previous studies, showing *H. pylori* was significantly more prevalent in apparently healthy people than celiac patients [15,16,18].

The protective effect of *H. pylori* on some allergic and atopic diseases and other inflammatory diseases could be considered as a starting point of the debates about the association between CD and *H. pylori* [22,23,40,48]. However, the findings were contradictory, and this association is not yet determined. There are different hypotheses and justification about the mechanism of interaction between *H. pylori* and CD. In the following paragraphs, we briefly discussed them.

Several studies had proposed the hygiene hypothesis, as an explanation for the protective role of *H. pylori* infection against CD. The hygiene theory is not limited to CD and has been proposed for other diseases such as asthma, irritable bowel disease (IBD), allergic rhinitis, and eczema [49,50]. It claims that lower exposure to some infectious agents in childhood leads to an increase in the risk of autoimmune diseases [18,22,23,40,51]. Some studies considered the increased prevalence of autoimmune diseases in Western environments alongside the increasing trend of health and cleanness, as an evidence for the validity of this theory [18,49,52]. Different mechanisms suggested for this hypothesis, but the role of each mechanism in the development of CD is unknown. One of them is T helper (Th) 1/Th2 deviation, which suggests that the presence of infection causes the balance of Th1/Th2 to move towards an immunosuppression state. T regulatory (Treg) lymphocytes might be induced or augmented (a phenomenon called bystander suppression) due to infection as an immunoregulation mechanism, which has associated with the Th1/Th2 deviation mechanism [49,51,53]. Another mechanism, antigenic competition/homeostasis, suggests that a strong immune response to infectious antigens reduces the immune response to ‘weak’ antigens, like allergens and autoantigens, seen in asthma and autoimmune diseases. Also, there are more suggested mechanisms for this hypothesis, such as non-antigenic ligands and gene-environment interaction, which may play a role in the hygiene hypothesis [49,51,53].

Another hypothesis is the protective role of *H. pylori* on the modification of gluten digested by proteases secretion, and increase of the pH of the gastric lumen, which reduces the immunogenicity of gluten [15,18,41,49,51,53]. Some studies have shown that consumption of protease (oral or protease of various microorganisms) leads to the digestion of gliadin (gluten protein). Of course, this method is not a substitute for a gluten-free diet, but it has been able to improve the lives of people who are sensitive to gluten (especially in cases of hidden gluten) to some extent. Although studies on *H. pylori* protease have not been fully performed, the results of studies on different proteases have shown a relationship in this regard [54,55].

Some studies support the idea that a specific virulence factor, such as CagA in *H. pylori*, may give the bacterium a protective role in CD. These studies detected less CagA^+^ *H. pylori* in patients with CD than the healthy control group, and also milder histological damage in CD patients infected with CagA^+^ *H. pylori* than CD patients infected with CagA^−^ *H. pylori*. This function of CagA may be due to the association with Treg cells in the immunoregulation mechanism [22,37,41,44].

One of the hypotheses that sought to explain the low prevalence of *H. pylori* in CD patients with severe histology came up with an interesting idea. It pointed out that patients with classic CD (10% – 40% of cases) who have classic symptoms such as chronic diarrhea, malabsorption, and abdominal pain may be misdiagnosed and treated for other GI diseases with similar symptoms, especially in developing countries [56,57]. Disease such as abdominal infection or small bowel bacterial overgrowth shows the same symptoms as classic CD, and it is more common [58,59]. Therefore, empirical antibiotic treatment can be prescribed for CD patients before the diagnosis of CD. This treatment may lead to the unintentional removal of *H. pylori* [42].

On the opposite side, some theories support the idea that *H. pylori* may be involved in the development of CD. The Second hit hypothesis proposed by Mooney et al. is an example of these theories. He mentioned that maybe *H. pylori* acts as an environmental trigger for CD, as observed in *Campylobacter* infection, in a study of the US military [40,52]. However, the mechanism of this act is not properly defined.

As *H. pylori* is a mucosa-associated bacterium, IgA type antibody is an essential part of the protection against this bacterium, at the primary level of the immune response in the gastrointestinal (GI) tract [60]. IgA deficiency is associated with CD that leads to false-negative IgA-based serology tests in CD patients [61,62]. The prevalence of IgA deficiency is 10-15 folds higher among CD patients than the general population, and recent studies show an increasing trend in its prevalence [63,64]. Although IgA deficiency does not have considerable prevalence, it may be one of the reasons for the high frequency of *H. pylori* in CD patients, which has not yet been addressed completely in the previous studies. It is suggested that future studies exclude CD patients with IgA deficiency, in order to remove the effect of this confounding factor.

For clearing the possible protective effect of the immune-suppressive mechanisms of *H. pylori* on CD, the first step is the specification of the chronological order of CD and *H. pylori* colonization. As there was not any cohort study on this subject, there is no strong evidence to show a cause and effect relationship between these two and a protective effect of *H. pylori* on the occurrence of CD. Therefore, conducting a cohort study should be considered as a top priority in further investigations of this subject.

Beyond our subgroup analyses, there are more interactive aspects of the pathophysiology of CD and pathogenesis of *H. pylori* that could help us to explore the causes of the heterogeneity. Both CD and *H. pylori* cause inflammation and immune response in the GI tract and also *H. pylori* could escape human immune defense via different mechanisms that suppress both innate and adaptive immune response [65]. *H. pylori* infection has a high inhibitory effect on leukocyte migration [66]. It also induces apoptosis of macrophages by alterations in the mitochondrial pathway [67]. Similar to *Mycobacterium Tuberculosis* (TB), interfering with lysosomal proteins let *H. pylori* to sustain intracellularly in macrophages [68]. *H. pylori* can evade host immune response via activating NF-κB and Wnt/β-catenin signaling pathway and disturbing metal ion homeostasis, due to the alteration of gene expression processes [69]. Nitric oxide (NO) is an important anti-microbial part of the innate immune response; production of arginase enzyme by *H. pylori* prevents the effect of NO on this organism and its lethal effect [70]. Recent studies reported that Vacuolating cytotoxin (VacA) of *H. pylori* has an immunomodulatory role by inhibiting integrin-linked kinase (ILK) [71] and interfering with the IL-2 signaling pathway in T-cells. VacA can block Ca^2+^ transportation and the activity of the Ca^2+^/calmodulin-dependent phosphatase calcineurin, therefore the organism can escape from the host immune response [72]. VacA-deficient *H. pylori* upregulates the expression of ILK and modulates the production of endothelial nitric oxygen synthase (eNOS), in comparison to isogenic VacA^+^ *H. pylori*. This process increases reactive oxygen species (ROS) in monocyte/macrophage-like U937 cells; therefore, it reduces the host immune reaction by damaging the immune cells [71]. VacA also intervenes with the antigen presentation of MHC class II [73] and it directly suppresses T-cells rather than antigen-presenting cells (APC) [74]. Another function of VacA is the interference with the expression of more than 100 related genes to immune evasion by changing their mRNA expression [75]. It is possible that these suppressive mechanisms produced by *H. pylori*, affect the autoimmune response to gluten and reduce the level of gluten enteropathy in CD. Although causality between *H. pylori* and CD is not detected yet, the coexistence of them in time and proximity of their pathogenic sites in the GI tract could be accounted as a basis for these interactions. The protective effect of *H. pylori* on CD is a production of a multivariate condition that immune suppressive mechanisms of *H. pylori* act as a counterbalance to the autoimmune response in CD, and reduce that to some degree. The *H. pylori*-related side of this condition depends on different factors such as ethnicity, chronicity of colonization, the quantitative scale of colonization, type of *H. pylori*, the serum level of antibody against *H. pylori*, *H. pylori* virulence factors, gastric ulcer, state of host immune system, and other variants that determine the range of immune suppressive mechanisms of *H. pylori*. The determinants of the CD-side of the condition could be marsh type, the severity of CD, the serum level of anti-tTg antibody, age at diagnosis, ethnicity, and other factors that show the severity of autoimmune response in CD patients. Considering the notion of a balance between immune suppressive mechanisms of *H. pylori* and autoimmune response in CD, it seems more probable to observe a protective effect against autoimmune response among patients who are infected with a type of *H. pylori* that produces the immune suppressive mechanisms and rises a good immune response, and on the other hand, the autoimmune response to gluten is not severe in these cases. These details were not investigated in previous studies and these could justify controversies about the protective role of *H. pylori* in CD patients.

In this systematic review and meta-analysis related to the association of *H. pylori* colonization and CD, our search strategy prepared a comprehensive source of articles that contains any study with related evidence. Subgroup and sensitivity analyses showed that two factors (sampling method and the quality of article) could be more effective and important for case-control studies and it is better to be addressed in future studies. In addition, the analyses showed that this protective effect of *H. pylori* on CD was more observed in Asian countries. After removing the studies with poor quality and non-appropriate sampling, the pooled OR was still statistically lower in a study from Asia, in comparison to both studies from Europe and America. However, heterogeneity was still more than 70% in this continent subgroup. Unfortunately, after removing the studies in sensitivity analysis, only one study from Asia remained, and the obtained results could not completely determine the source of heterogeneity. Here, we suggest conducting well-designed studies in Asia, especially in the East Asian population to clarify the role of ethnicity in the association of *H. pylori* colonization and CD.

The present study had some limitations. First, just a few numbers of the selected articles had a good-quality method and many of them did not consider many confounding factors in the design of the study such as March type, the severity of celiac, *H. pylori* virulence, and IgA deficiency that could affect the results. Second, based on the results, the heterogeneity of outcomes was high, and the subgroup analysis could not recognize the source of it completely. Third, most of the studies were conducted in Western countries, especially in Turkey and Italy. There was no article from Africa. Moreover, only three articles reported the situation in Asia. Studying other ethnicities can be a suitable target for further investigations.

## Conclusion

The meta-analysis of 24 clinical studies included 5241 CD patients from both children and adult age-groups, showed that *H. pylori* was associated negatively with CD. However, there is not enough evidence to indicate a strong protective effect of *H. pylori* on CD. The main obstacle on the way of placing the role of *H. pylori* in CD under scrutiny is the shortage of accurate details in the clinical studies. To clear the relationship between *H. pylori* and CD, it is suggested to implement cohort studies and also further primary studies with good methodology. Extending the scope of study about this subject to the molecular context may prepare more answers to the current controversies.

## Specific author contributions

Guarantor of the article: SB and HS-A. Development of study concept and design: SB, HS-A, FIA, and AN. Acquisition of data: SB, ZN, NG, SAF, AA, and AN. Analysis and interpretation of data: HS-A, SB, AS, and FIA. Statistical analysis: AS. Drafting of the manuscript: HS-A, SB, and FIA. Project administration: HS-A, and SB. Funding acquisition: SB, and ZN. Critical revision of the manuscript for important intellectual content: all authors. Comment on drafts of the paper: HS-A, SB; and approve the final draft of the manuscript: all authors.

